# Oral–Gut Microbial Axis in Inflammatory Bowel Disease and Primary Sclerosing Cholangitis

**DOI:** 10.64898/2026.07.15.738466

**Authors:** Shanlin Ke, Zihan Zhou, Xiaole Yin, Yiyan Yang, Zheng Sun, Fernanda Quevedo, Nadine Javier, Malav Dave, Katherine Wu, Shaikh Danish Mahmood, Yang-Yu Liu, Joshua R. Korzenik

**Affiliations:** Channing Division of Network Medicine, Department of Medicine, Brigham and Women’s Hospital and Harvard Medical School, Boston, MA 02115, USA; Division of Gastroenterology, Hepatology and Nutrition, Department of Internal Medicine, College of Medicine, The Ohio State University Wexner Medical Center, Columbus, OH 43210, USA; The James Comprehensive Cancer Center, Ohio State University Wexner Medical Center, Columbus, OH 43210, USA; School of Civil and Environmental Engineering, Nanyang Technological University, Singapore 639798; Division of Gastroenterology, Hepatology and Endoscopy, Brigham and Women’s Hospital, Harvard Medical School, Boston, MA 02115, USA

**Author notes:** Correspondence should be addressed to J.R.K. and Y.-Y.L.

## Abstract

The oral cavity is increasingly recognized as a reservoir of microbes that can translocate to and influence the gut microbiome. This oral–gut microbial axis may contribute to the pathogenesis of chronic gastrointestinal and hepatobiliary disorders, including inflammatory bowel disease (IBD) and its comorbidity, primary sclerosing cholangitis (PSC). To investigate the oral–gut microbial axis in IBD and PSC, we enrolled 191 participants spanning Crohn’s disease (CD), ulcerative colitis (UC), CD with PSC (CD-PSC), UC with PSC (UC-PSC), and healthy controls. We generated and analyzed the whole-metagenome shotgun sequencing data from saliva, tongue swabs, and fecal samples. Across oral and gut niches, we identified multiple microbial species differentially enriched in participants with UC compared with healthy controls, including *Fusobacterium nucleatum* and *Gemella sanguinis* in saliva, *Catonella massiliensis* and *Tannerella serpentiformis* on the tongue, and *G. sanguinis* and *Sellimonas intestinalis* in feces. Paired oral-gut analyses revealed 15 potential oral-origin species enriched in fecal samples; notably, *G. sanguinis* and *Veillonella rogosae* were consistently enriched in participants with UC. These findings were further supported by an external validation dataset comprising 1,716 gut metagenomes from five independent IBD cohorts, highlighting the reproducibility of oral microbial signatures. In exploratory analyses, smoking history was associated with shifts in the oral microbiome toward a UC-like state in healthy controls, suggesting a possible environmental modifier of the oral-gut microbial axis. Collectively, our study identifies distinct oral microbial signatures and their potential translocation to the gut in IBD and PSC, underscoring the role of the oral– gut microbial axis in disease progression.

## INTRODUCTION

The human microbiota is not only represented by the microbes from the gastrointestinal tract, but also those resident in other body sites, including the skin, oral cavity, lung, and urinary tract^1^. Those microbial communities are not entirely isolated from one another; they interact and influence each other through various pathways. Among these, the oral-gut microbial axis has gained increasing attention.

The oral cavity, as a primary gateway to the gastrointestinal tract^2^, harbors a distinct microbial community shaped by various factors such as pH, oxygen levels, nutrient availability, and the host immune response^3^. Accumulating evidence indicates that resident oral microbes can enter the gut through swallowing, aspiration, or translocation^4^. This translocation can occur even in healthy individuals^5^. Under certain conditions, the translocation of oral microbes may contribute to or exacerbate gastrointestinal diseases^6^.

Recent studies have highlighted the importance of the gut microbiome in the pathogenesis of inflammatory bowel disease (IBD)^7–11^ and its comorbidity with primary sclerosing cholangitis (PSC)^12^. IBD, primarily Crohn’s disease (CD) and ulcerative colitis (UC), cause inflammation and damage to the digestive tract, resulting in diarrhea, abdominal pain, and weight loss. PSC is a chronic and progressive cholestatic disease characterized by peribiliary inflammation and fibrosis^13–16^. An estimated 60-80% of individuals with PSC are also diagnosed with IBD^17^. However, the involvement of oral microbiome in the pathogenesis of IBD and PSC, particularly via oral–gut translocation, remains poorly understood.

Cigarette smoking is an important environmental factor that can alter both oral and gut microbial communities and has implications for disease development^18^. Interestingly, previous studies have suggested that smoking has opposing effects in UC and CD^19,20^. In general, smoking is associated with a reduced risk of developing UC, while former smokers may have an increased risk^21^ and in some cases quitting smoking may even worsen UC symptoms^22^. By contrast, smoking is widely recognized to worsen the course of CD, leading to more severe disease and a higher rate of complications^23,24^. The factors underlying these opposing effects remain largely unclear. One possible mechanism is that smoking may influence the oral microbiome through the induction of anaerobic conditions, suppression of local immune defenses, shifts in salivary pH, and the antimicrobial effects of toxic compounds present in cigarette smoke^25^. These changes may disrupt oral and gut microbial communities in ways that contribute to the pathogenesis of UC and CD differentially. Recently, it was demonstrated that smoking is associated with increased oral bacteria in the colonic mucosa and affects the gut immune system^26^.

In the present study, we leveraged a well-characterized cohort ORCA (Oral Microbiome Assessment in PSC) to investigate the distinct oral microbial signature related to IBD and PSC, the impact of smoking on the oral microbiome, and its potential contribution to the pathogenesis of IBD and PSC (**Fig. 1a**). We performed whole-metagenome shotgun (WMS) sequencing of microbiome samples from the oral cavity (saliva and tongue swab) and gut (feces) to examine the connections between the oral and gut microbiomes, providing new insights into their potential roles in the pathogenesis of IBD and PSC.

**Fig. 1.**
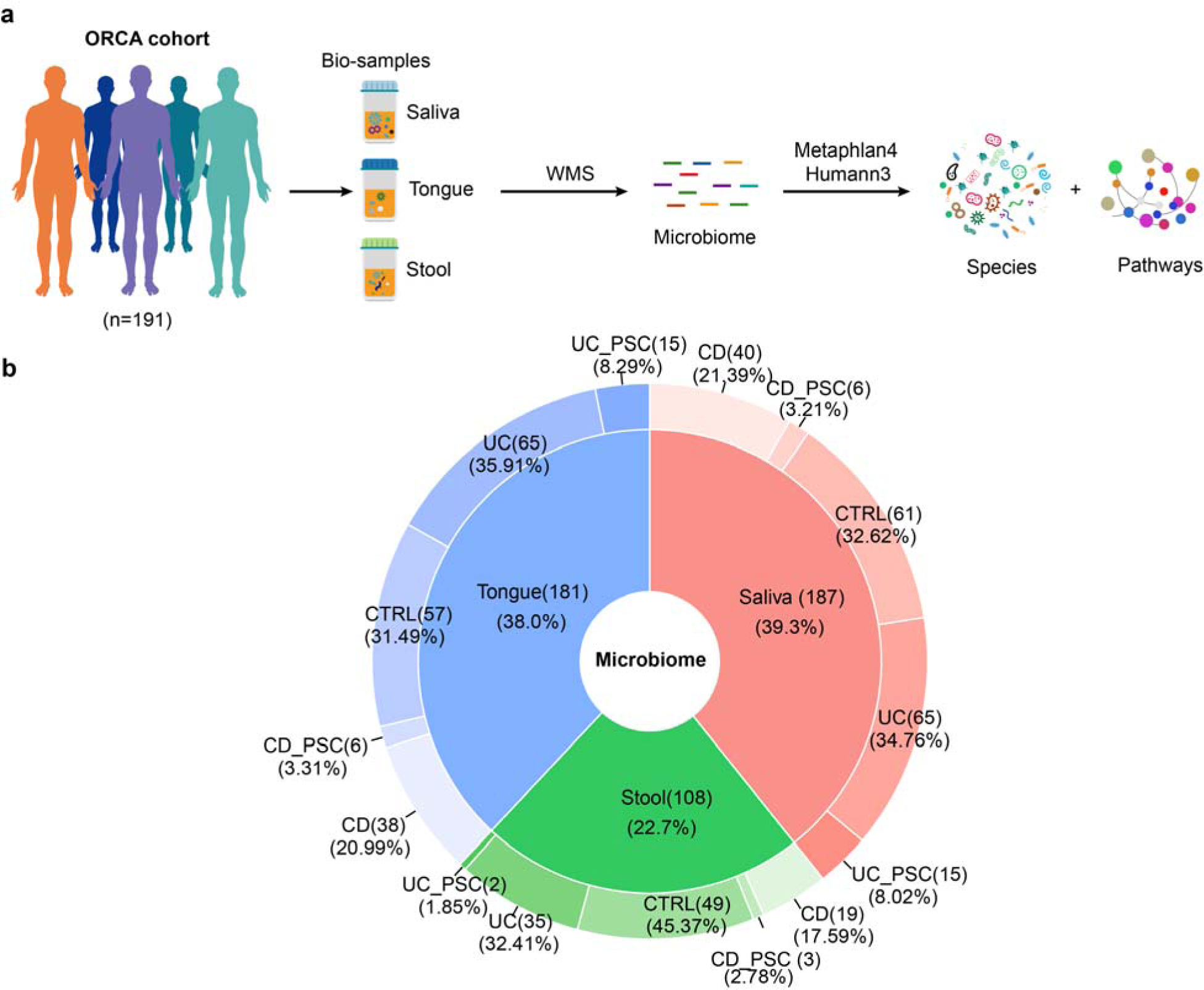
Workflow and the study cohort. **a**. A conceptual framework of this study. A total of 191 eligible participants were enrolled, including healthy controls, CD alone, CD-PSC, UC alone, and UC-PSC. Samples were collected from three body sites: saliva, tongue swab, and feces. All sample types underwent shotgun metagenomic sequencing. **b**. Sample distribution of microbiome among different body sites and phenotypic groups.

## RESULTS

### Study cohort

In the ORCA cohort, we enrolled 191 participants, including individuals with CD alone (n=42), UC alone (n=65), CD with PSC (CD-PSC, n=6), UC with PSC (UC-PSC, n=16), and healthy controls (n=62). The healthy control group included current smokers (n=22), former smokers (n=20) and never smokers (n=20). We collected saliva, tongue swabs, and fecal samples from those individuals and performed WMS sequencing (**Fig. 1a, Methods**). In total, we have WMS sequencing data from 201 saliva samples (from 187 individuals), 195 tongue samples (from 181 individuals), and 109 stool samples (from 108 individuals) (**Fig. 1b**). For 103 individuals, we have the WMS data from all three sample types.

### Characterization of the microbiome in different body sites

For the WMS sequencing data, we applied MetaPhlan4^27^ for taxonomic profiling. We found that fecal samples contained a significantly larger number of microbial species than saliva and tongue swab samples (**Fig. 2a**). These results indicate the complexity of the gut compared to the oral cavity in terms of microbiome diversity. Consistent with previous findings, the microbial composition of the fecal microbiome was quite distinct from that of the oral microbiome, while saliva and tongue samples shared similar microbial compositions (**Fig. 2b**).

**Fig. 2.**
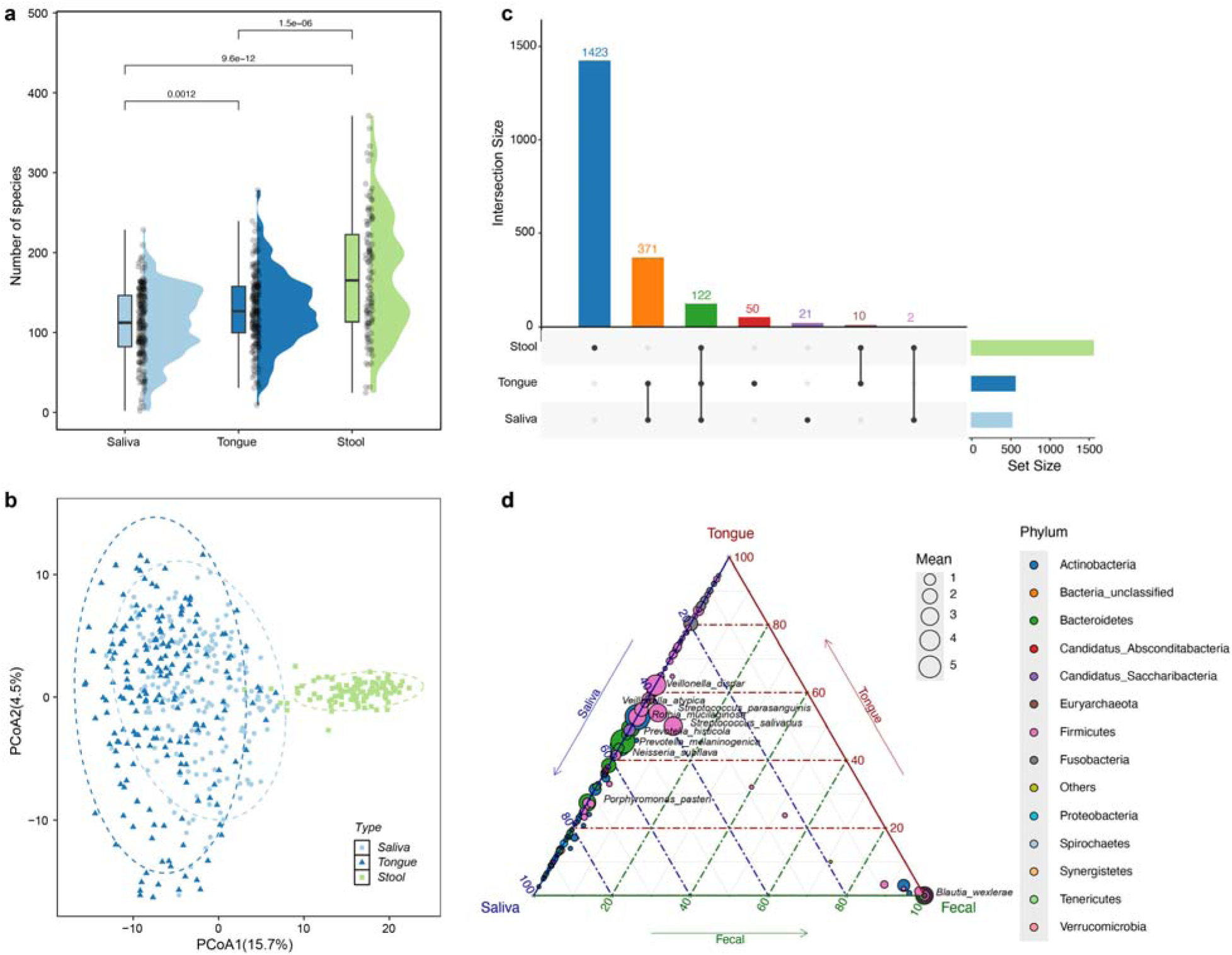
Distinct oral and gut microbiome. (**a**), Number of microbial species identified from different body sites. (**b**), Principal Coordinates Analysis (PCoA) plot of the microbiome based on robust Aitchison distance on samples from different body sites. (**c)**, The UpSet plot shows the intersections of identified species across different body sites, respectively. (**d**), Ternary plots of per-site mean relative abundances of species.

Among the species identified from the three body sites, 122 shared species were identified (**Fig. 2c**). We found multiple species were shared by saliva and tongue swabs and have similar relative abundances, such as *Neisseria subflava*, *Prevotella melaninogenica*, *Rothia mucilaginosa*, and *Streptococcus parasanguinis* (**Fig. 2d**). However, multiple species also demonstrated significant differences in abundance between saliva and tongue samples among distinct phenotypic groups (**Fig. S1**). For example, *Gemella* species (e.g., *G. sanguinis*, *G. morbillorum*, and *G. haemolysans*), *Fusobacterium* (e.g., *F. pseudoperiodonticum* and *F. nucleatum*), *Leptotrichia* species (e.g., *L. hongkongensis and L. wadei*), and *Solobacterium species* (e.g., *S. moorei and S. SGB6829*) were more abundant in tongue swab samples, whereas *Prevotella species* (e.g., *P. maculosa*, *P. nigrescens*, and *P. oris*) and *Porphyromonas* species (e.g., *P. pasteri*, *P. endodontalis*, and *P. catoniae*) were more enriched in saliva samples. Similarly, differences were also observed at the level of functional capacity between saliva and tongue samples (**Fig. S2**). Compared to oral samples, species like *Blautia wexlerae* were more abundant in fecal samples (**Fig. 2d**).

### Characterization of microbiome alteration in IBD and PSC

We measured the alpha diversity using the Shannon index of the microbiome samples of different disease groups and healthy controls. For saliva samples, no significant differences of alpha diversity were found between disease groups and controls (**Fig. 3a**). For tongue swab samples, the UC group exhibited a significantly higher Shannon index than healthy controls (**Fig. 3b**). For fecal samples, the UC-PSC group showed a significantly lower Shannon index than healthy controls (**Fig. 3c**).

**Fig. 3.**
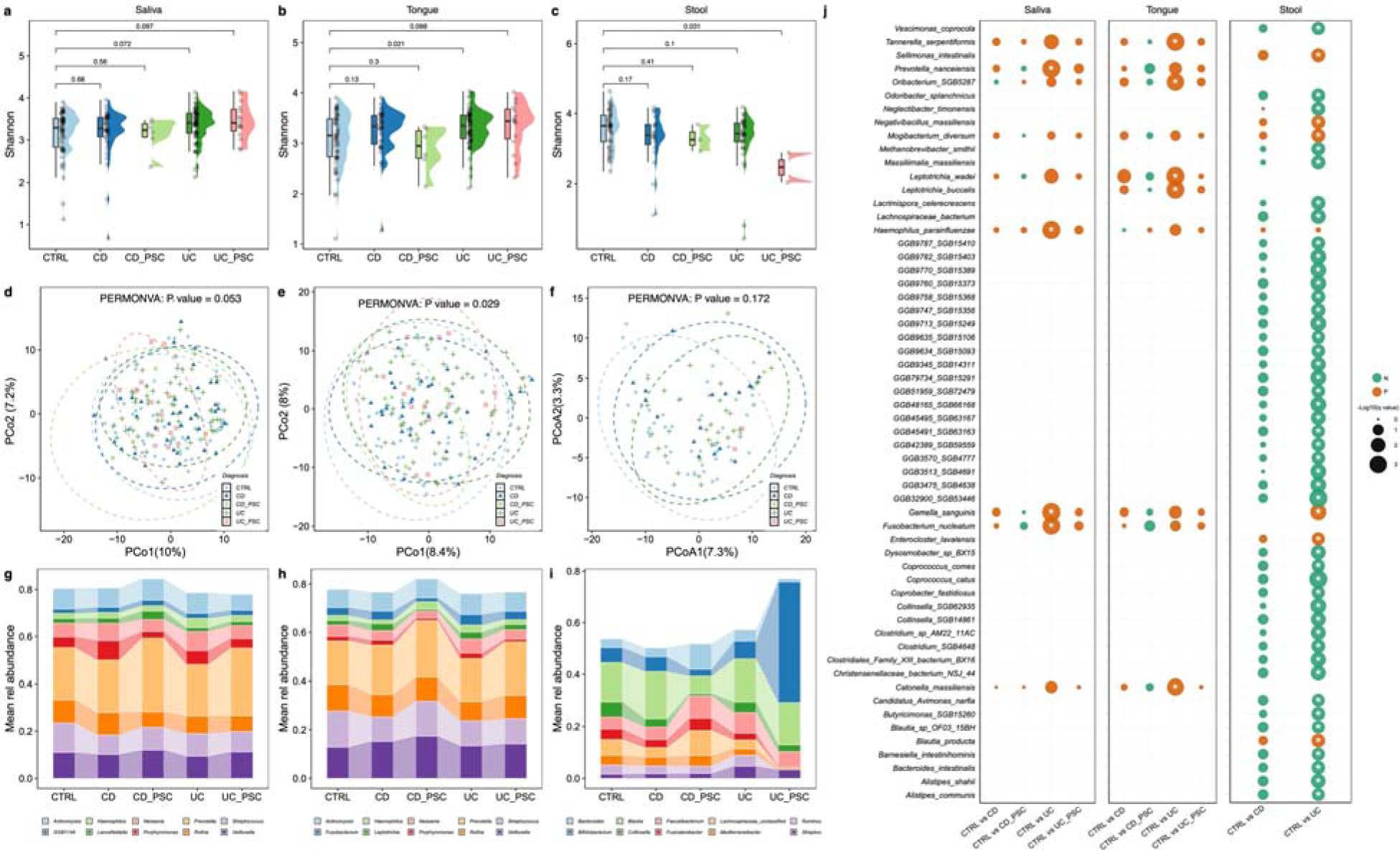
IBD and PSC-related differences of the human oral and gut microbiome. Violin box plots of alpha diversity of taxonomic profiles at the species level using Shannon index in different groups from saliva (**a**), tongue (**b**), and stool (**c**). P-values were calculated by two-sided Wilcoxon–Mann–Whitney test. PCoA of microbiome samples from different groups at the species level based on the robust Aitchison distance (**d**-**f**). The abundance of the top 10 main genera identified from different groups in saliva (**g**), tongue (**h**), and stool (**i**). The associations between IBD and PSC and microbial species were identified using MaAsLin2 (**j**). Significant species-level features from each disease group are summarized (from left to right are saliva, tongue, and stool). Only statistically significant associations with q value ≤ 0.25 (Benjamini Hochberg adjusted P value) are labeled with an asterisk. The size of each dot represents the - log10 (q value). The color of each dot represents the valence of the association: green=negative, enriched in controls, orange =positive, absent in controls.

As for the beta diversity, Principal Coordinate Analysis (PCoA) based on the robust Aitchison distance revealed that only microbial compositions of tongue swab samples significantly differed among disease groups (**Fig. 3d-f**, PERMANOVA p-value = 0.029).

At the genus level, we found that CD-PSC group carried more *Prevotella* in their saliva and tongue samples than other disease groups (**Fig. 3g-h**), and UC-PSC patients carried significantly more *Bifidobacterium* in their fecal samples (**Fig. 3i**).

We next used general linear models with age and gender adjusted, as implemented in microbiome multivariable associations with linear models (MaAsLin2)^28^, to identify differentially abundant species in CD, CD-PSC, UC, and UC-PSC, with respect to healthy controls. We identified multiple species differentially abundant in UC group compared with controls (false discovery rate (FDR) < 0.25) in the three sample types. In saliva, we found multiple species enriched in participants with UC compared to healthy controls, such as *F. nucleatum*, *Gemella sanguinis*, *Haemophilus parainfluenzae*, and *Prevotella nanceiensis* (**Fig. 3j**). We identified five tongue species that were enriched in UC group, including *Catonella massiliensis*, *Leptotrichia buccalis*, *Leptotrichia wadei*, *Oribacterium SGB5287*, and *T. serpentiformis.* In fecal samples, 6 enriched species (e.g., *Blautia producta*, *G. sanguinis,* and *Mogibacterium diversum*) and 44 depleted species (e.g., *Alistipes shahii*, *Barnesiella intestinihominis*, and *Coprococcus catus)* were found in participants with UC compared with healthy controls. Combining CD-PSC and UC-PSC together as IBD-PSC, we found that *Neisseria bacilliformis* was enriched in both saliva and tongue samples from patients with IBD-PSC compared to those with IBD alone, while *Porphyromonas pasteri* was depleted in saliva in IBD-PSC. When comparing CD-PSC to UC-PSC, *Blautia luti* was depleted in fecal samples and *Selenomonas SGB5900* was depleted in saliva (**Fig. S3**).

### Identification of oral bacteria in human fecal samples

To determine whether there are any translocation events that may be related to IBD or PSC, we first calculated species sharing rates between body sites in 103 participants for whom we had microbiome data from all three sample types: saliva, tongue swab, and fecal samples. For each participant, we computed the proportion of shared species between two sample types by dividing the number of overlapping species by the total nonredundant species in the union of both sample types. Consistent with the findings from a previous study^29^ and comparison at the population level (**Fig. 2c**), we found that saliva and tongue microbiome samples shared more species with each other than with stool samples (**Fig. 4a-c**). In participants with CD, their saliva samples shared significantly more species with tongue swab samples compared to controls (**Fig. 4a**). More interestingly, we found that participants with UC shared significantly more species between their oral microbiomes and fecal microbiome compared to controls (**Fig. 4b-c**). This trend was not observed in other disease groups, i.e., CD and IBD-PSC. This finding suggests that oral microbial species may play a role in UC via translocation.

**Fig. 4.**
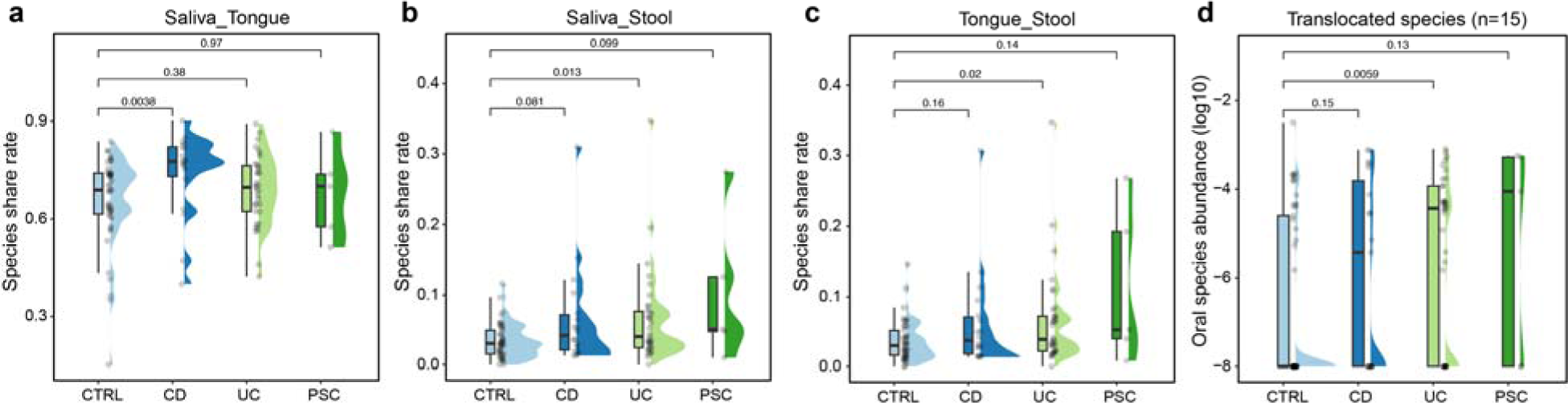
Identification of oral-origin translocated species in the human fecal samples. Species sharing rate in pairs of saliva-tongue (**a**), saliva-stool (**b**), and tongue-stool (**c**) across different groups. The total abundance of 15 oral-origin translocation species in stool samples across different groups (**d**).

To quantify oral bacteria in human fecal samples, we adopted the method developed in a previous study^26^ to compare the relative abundance and prevalence of oral and fecal samples (see **Methods**). We identified a total of 15 potentially oral-origin species, including *Actinobaculum sp oral taxon 183*, *Actinomyces dentalis, Actinomyces SGB17163, Actinomyces sp oral taxon 448*, *Aggregatibacter segnis, Corynebacterium durum*, *Gemella sanguinis, GGB12785 SGB19823*, *Parvimonas micra*, *Scardovia wiggsiae*, *Schaalia odontolytica*, *Streptococcus cristatus*, *Streptococcus rubneri*, *Veillonella rogosae*, and *Veillonella tobetsuensis*. Notably, the total relative abundance of these 15 species was significantly higher in the fecal samples of UC patients than in healthy controls (**Fig.4d**). A similar trend was not observed for other disease groups, i.e., CD and PSC. These findings are consistent with the findings from the species sharing rates (**Fig. 4b,c**). Among these 15 species, nine of them (*A. dentalis, A. segnis, C. durum, G. sanguinis, P. micra, S. wiggsiae, S. odontolytica, S. cristatus and V. rogosae*) were listed in eHOMD^30^, a database that provides comprehensive curated information on bacteria in the human mouth and aerodigestive tract, including the pharynx, nasal passages, sinuses and esophagus. The consistent findings from Fig.4b,c and Fig.4d imply that oral-gut microbial translocation very likely occurs in UC, rather than other disease groups, i.e., CD and PSC.

### Effect of smoking on the human oral and gut microbiome in healthy controls

To explore if the human microbiome may be involved in the opposing effects of smoking on UC and CD, we first compared the microbial compositions and functional profiles of healthy controls with different smoking statuses: current smokers, former smokers, and never smokers. We found that smoking status was significantly associated with the overall microbial structure of tongue samples, but not saliva or stool samples (**Fig. 5a-c**).

**Fig. 5.**
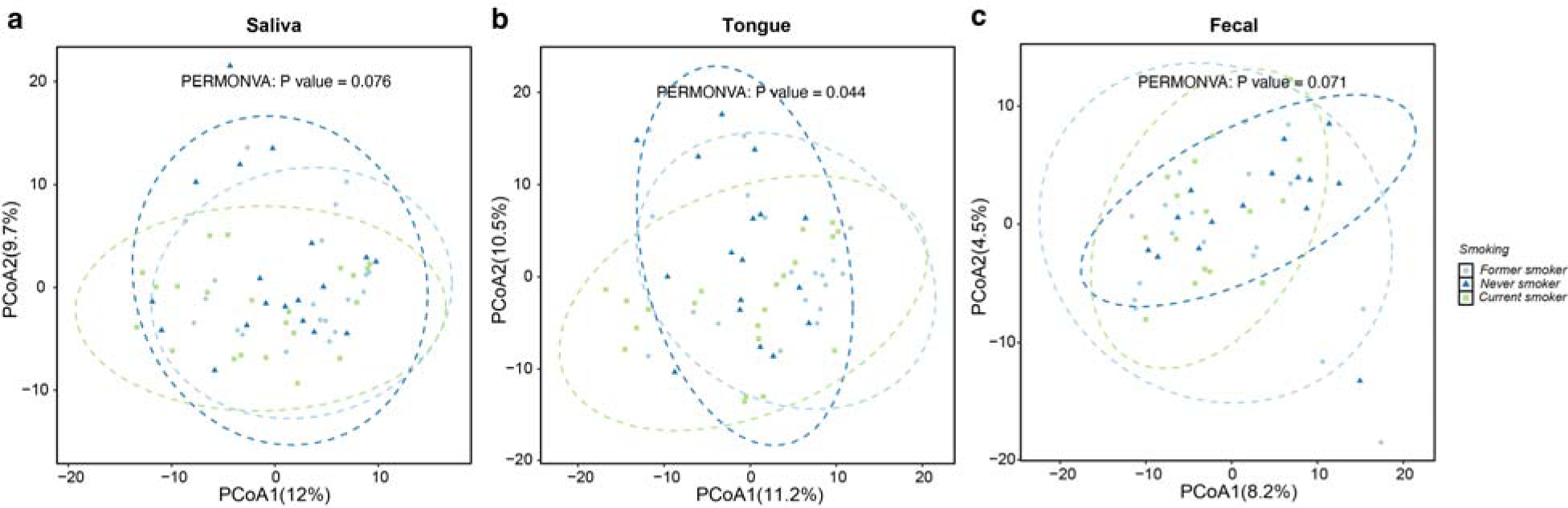
Effect of smoking on the human oral and gut microbiome in healthy controls. (a)-(c). PCoA plot of microbiome samples from different smoking statuses at the species level based on the robust Aitchison distance.

In saliva, we identified 20 differentially abundant species between current and former smokers, including *Actinomyces graevenitzii*, *L. wadei*, and *Prevotella pallens* (**Fig. S4**). However, no significant differences in species abundance were observed between current and never smokers, or between former and never smokers. In tongue samples, we found two species (*Candidatus Saccharibacteria unclassified SGB95587* and *Cardiobacterium hominis*) differentially abundant between current and never smokers, while *P. pallens* was enriched in current smokers compared to former smokers (**Fig. S5**).

At the functional pathway level, smoking status was not significantly associated with the overall functional profile (**Fig. S6**). However, we identified multiple arginine biosynthesis related pathways (e.g., ARGSYNBSUB-PWY, ARGSYN-PWY, PWY-5154, and PWY-7400) that were significantly downregulated in the saliva sample of current smokers compared to former smokers (**Fig. S7a**). A similar reduction in these pathways was observed in the tongue swab samples of current smokers compared to former smokers. Additionally, ARGSYN-PWY and PWY-7400 were decreased in former smokers compared to never smokers (**Fig. S7b**).

### Smoking history shapes oral microbiome similarity to participants with UC

In the ORCA cohort, unfortunately, no current smokers were enrolled among participants with UC, and only one current smoker with CD was enrolled. The healthy control group did include individuals with varying smoking statuses. To test how smoking is associated with the oral and gut microbiome in IBD, we stratified the healthy controls into three different subgroups based on their smoking status (never smoker, former smoker, and current smoker, denoted as HC_ns_, HC_fs_, and HC_cs_) and then calculated the microbiome distance between participants with UC/CD and the three subgroups of healthy controls. Interestingly, we found that the oral microbiome distance between UC and HC_cs_ is significantly higher than that between UC and HC_ns_ or HC_fs_, regardless of the sample types: saliva or tongue swab (**Fig. 6a-b**). Yet, the trend is opposite for stool samples. The stool microbiome distance between UC and HC_cs_ is significantly lower than that between UC and HC_ns_ or HC_fs_ (**Fig. 6c**), indicating sample types can be very critical to identify the positive or negative effects of smoking on UC. However, we did not observe similar pattern in patients with CD (**Fig. S8**).

**Fig. 6.**
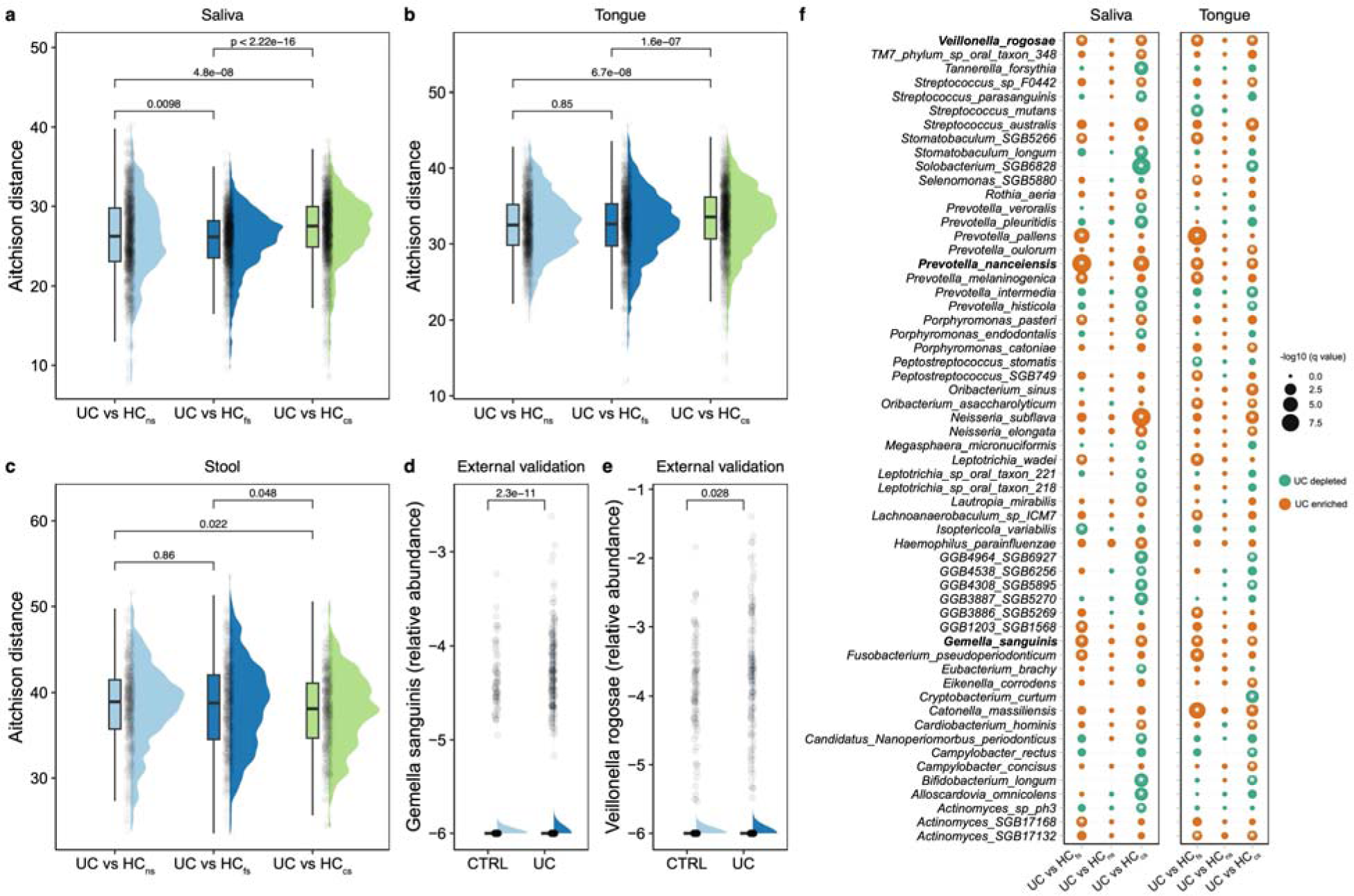
Smoking history shapes oral microbiome similarity to participants with UC. Microbial differences between the smoking group and UC group quantified by robust Aitchison distance in saliva (**a**), tongue swab (**b**), and stool (**c**) samples. *P*-values were calculated by two-sided Mann–Whitney test. External validation of *G. sanguinis* (**d**) and *V. rogosae* (**e**) in the gut samples of UC across five independent cohorts. P-values were calculated by the one-sided Mann–Whitney test. (**f**) The differential abundant species between the smoking group and UC group were identified using MaAsLin2. Significant species-level features from each comparison are summarized (from left to right are saliva and tongue). Only statistically significant associations with *q* value ≤ 0.25 (Benjamini Hochberg adjusted *P* value) are labeled with an asterisk. The size of each dot represents the -log10 (*q* value). The color of each dot represents the valence of the association: green=negative, depleted in UC group, orange =positive, enriched in UC group. HC_ns_: Never smoker in Control group; HC_fs_: Former smoker in Control group; HC_cs_: Current smoker in Control group.

We further compared oral microbial compositions to identify specific species driving the observed difference between participants with UC and the three subgroups of healthy controls. Notably, three specific species (i.e., *G. sanguinis*, *P. nanceiensis*, and *V. rogosae*) were significantly enriched in UC group compared to HC_cs_ and HC_fs_ in both saliva and tongue samples (**Fig. 6f**). However, these species were not differentially abundant between UC and HC_ns_. Interestingly, both *G. sanguinis* and *V. rogosae* have been identified as potential oral-origin species detected in fecal samples (**Fig.6e**), suggesting that smoking may influence the translocation of microbes from the oral cavity to the gut.

To further validate the translocation of *G. sanguinis* and *V. rogosae* from the oral cavity to the gut, we analyzed 1,716 gut metagenomic samples from healthy controls and participants with UC in an external validation dataset (consisting of five independent cohorts^10,31–35)^. Importantly, we found that both *G. sanguinis* and *V. rogosae* were more abundant in the gut microbiome of participants with UC compared to healthy controls (**Fig. 6d-e**).

## DISCUSSION

This study comprehensively measures associations between oral and gut microbiome in a well-characterized cohort of IBD and PSC. Our results provide critical insights into the interplay between the oral and gut microbiomes and their associated metabolic profiles in IBD and PSC. Through systematic analysis of saliva, tongue swabs, and fecal samples, we identified significant alterations in microbial composition, diversity, and functional capacity, along with evidence of oral microbial translocation in UC and the potential modulatory role of smoking. These results shed light on novel microbial and metabolic mechanisms that may contribute to disease pathogenesis and progression of UC.

Over the past decades, the role of the human gut microbiome in IBD has been widely explored^7,9,36,37^. In contrast, the potential contribution of microbes from other body sites, such as the oral cavity, has received comparatively less attention. Previous studies have demonstrated that IBD patients are associated with a significantly higher prevalence of periodontitis^38,39^, suggesting the potential role of the oral microbiome in IBD. Among the different phenotypic groups, we found in this study that UC patients were associated with more oral species, such as *F. nucleatum*, *P. nanceiensis*, *T. serpentiformis*, and *G. sanguinis*. As a periodontal pathogen, *F. nucleatum* has been shown to exacerbate the severity and microbial dysbiosis in DSS-induced colitis mice^40^, and to promote gut inflammation and epithelial barrier dysfunction, thereby aggravating UC^41^. *T. serpentiformis*, commonly found in oral biofilms, has been linked to periodontitis^42^. Additionally, *G. sanguinis*, previously implicated in infective endocarditis^43,44^, was enriched in both oral and gut microbiome of UC patients. Meanwhile, we identified more species that were absent in UC patients in the gut samples, for example, *A. shahii*, *B. intestinihominis*, and *C. catus*. These species have also previously been reported to be depleted in UC patients^45^ or associated with beneficial effects in DSS-induced colitis^46,47^.

Although physical and chemical barriers like bile and stomach acid typically separate the oral and gut microbiomes, the disruption of the oral-intestinal barrier can enable microbial translocation between the two sites and contribute to diseases. Leveraging the paired oral (saliva and tongue) and fecal samples, we found UC patients shared more species between oral and fecal samples, suggesting increased oral-gut microbial translocation or barrier disruption. Importantly, the 15 identified oral-origin species were indeed significantly more abundant in UC patients. These findings align with a previous study of 2,932 paired oral (collected from several oral cavity sites) and fecal samples with 16S rRNA gene sequencing from 223 healthy individuals in the Human Microbiome Project, which identified several oral-origin genera, including *Actinomyces*, *Aggregatibacter*, *Corynebacterium*, *Parvimonas*, *Streptococcus*, and *Veillonella* in human feces^48^. Notably, most of the 15 species we identified belonged to these genera. Additionally, *G. sanguinis* was identified as an oral-origin bacteria, consistent with our observation of its enrichment in both oral and gut samples from UC patients.

In our cohort, although current smokers were not included among the UC patients, the control group included individuals with varying smoking statuses. By comparing UC patients with controls stratified by smoking status, we found that the oral microbiomes of current and former smokers of healthy controls differed more from those of UC patients than did those of never smokers. Interestingly, this distinction was primarily driven by two oral-origin bacteria (*G. sanguinis* and *V. rogosae*), suggesting that smoking status may influence oral-gut translocations. Notably, these two species were present at similar levels in never smokers of healthy controls and UC patients, leading us to hypothesize that smoking may exert a protective effect in UC by reducing oral-gut microbial translocation. Furthermore, we confirmed that *G. sanguinis* and *V. rogosae* were significantly enriched in the UC patients across 1,716 external gut metagenomic samples, supporting their translocation from the oral cavity to the gut and their potential role in UC pathogenesis. However, more research is needed to determine the underlying mechanisms behind these findings. These smoking-related findings should be regarded as hypothesis-generating rather than conclusive, given the small PSC subgroups (e.g., CD-PSC, n = 6), the absence of current smokers among participants with UC and the use of different taxonomic profilers for the ORCA discovery cohort (MetaPhlAn4) and the external validation cohorts (Phanta), which precludes direct quantitative comparison across datasets.

Importantly, we observed that oral–gut microbial translocation signals were predominantly detected in UC but not in CD, and that smoking-associated differences in oral microbiome composition were also primarily evident in UC rather than CD. This subtype-specific pattern is notable given that previous epidemiological studies have shown that smoking is differentially associated with IBD subtypes, being protective in UC but a risk factor for CD^20^. Although our findings do not directly address causality, they suggest that smoking may be associated with modulation of oral–gut microbial interactions in a disease-specific manner, potentially reflecting distinct host– environment–microbiome relationships between UC and CD.

Previous studies have also investigated the oral microbiome in PSC and provide important insights on host-microbiota associations in PSC. Lapidot et al. (17 PSC, 18 PSC-IBD, and 30 controls)^49^ found that adult PSC patients exhibit significant alterations in both salivary and fecal microbiomes, with enrichment of oral-associated genera such as *Veillonella* and *Streptococcus* in both fecal and saliva of patients with PSC, suggesting oral–gut microbial overlap. Iwasawa et al. (24 PSC, 16 UC, and 24 healthy controls)^50^ found that pediatric-onset PSC (mean age ∼12.13 years) showed distinct salivary beta-diversity compared with UC and healthy controls, with decreased *Rothia* and *Haemophilus* versus healthy controls and decreased *Haemophilus* but increased *Oribacterium* versus UC, despite no significant alpha-diversity differences. Collectively, these studies support that PSC is associated with oral microbiome alterations, although differences in findings across studies may be attributed to variations in sample size, population characteristics (adult vs pediatric cohorts), sampling strategies (saliva alone vs paired oral–gut sampling), and sequencing resolution (16S rRNA gene profiling in prior studies versus shotgun metagenomics in our study).

The translocation of oral bacteria to the gut has several important implications in addition to understanding the persistent dysbiosis characteristic of UC, CD and PSC. There remains strong interest in manipulating the intestinal bacteria through fecal transplant or other means^51–54^. These efforts may be less effective than hoped for as the oral microbiome may serve as a reservoir to perpetuate the dysbiosis evident in these diseases. The recurrent recolonization from the mouth to the gut may contribute to frustrating these efforts to renovate the gut microbiome more substantially or for a longer duration.

In conclusion, our study contributes to the understanding of the oral microbiome in IBD and highlights the potential role of smoking in modulating disease risk in UC. These insights have important implications for understanding the mechanisms underlying oral-gut translocation, which may potentially be targeted in preventive or therapeutic strategies for IBD.

## Methods

### ORCA participants enrollment

Patients were enrolled in the BWH Crohn’s and Colitis Center who were 18 years or older with a confirmed diagnosis of PSC or IBD for over 3 months. They would need to be on stable IBD meds for at least 4 weeks prior to enrollment. Patients were excluded if they (1) had a dental procedure within 30 days of enrollment; (2) used antibiotics within 6 weeks of sample collection; or (3) chronically used acid-suppressing medication (proton pump inhibitor, H2-receptor antagonist, or antacids) or nitroglycerin/nitrate preparations, defined as use ≥ 3 times per week for ≥ 3 months. Subjects were required to be fasting for at least 3 hours prior to the oral sample collection and collections were all done in the morning before 1 PM. The study was reviewed and approved by the Institutional Review Boards (IRB) of Mass General Brigham (MGB). Written informed consent was obtained from all participants prior to sample collection.

### Sample collection

Saliva was collected in an OMR-120 dnagenotek tube for tongue swab and an OMR-501 dnagenotek tube for saliva global microbiome collection. Aliquots were made immediately at collection into cryovials for metabolomics. Samples were flash frozen within 10 minutes of collection and stored at -80°. The tongue swab was collected from the rear dorsal area of the tongue prior to saliva collection and stored at -80°. Fecal samples were collected prior to the start of a colonoscopy prep or if not preparing for a colonoscopy within 2 days of the saliva collection and frozen at home if not brought in the day of collection.

### DNA extraction and metagenome sequencing

Oral and stool samples were stored at -80 ℃ before sending them to Diversigen for DNA extraction, whole-genome shotgun library preparation, and Illumina sequencing. The metagenomics sequencing targeted approximately 5 Gb of sequences per sample, utilizing 151 base pair paired end reads.

### Microbiome taxonomic and functional potential profiling

We performed quality control on the raw metagenomics sequencing data by initially discarding low-quality reads. Subsequently, we removed reads belonging to the human genome by mapping the data to the human reference genome with KneadData (https://huttenhower.sph.harvard.edu/kneaddata/). Microbial taxonomic profiling was performed using MetaPhlAn4^27^, which uses a library of clade-specific markers to provide panmicrobial (bacterial, archaeal, viral, and eukaryotic) profiling. We then performed functional profiling for metagenomes by applying HUMAnN3^55^, which maps DNA reads to a customized database of functionally annotated pan-genomes.

### Identification of oral species in fecal samples

An oral-origin species in a fecal sample is defined as one that meets the following four criteria^48^: 1) The species’ average relative abundance across all oral cavity samples exceeds *θ_a_*; 2) The species’ average relative abundance across all fecal samples does not exceed *θ_a_* but large than 0; 3) The species’ prevalence across all oral cavity samples exceeds *θ_p_* ; and 4) The species’ prevalence across all fecal samples does not exceed but is larger than *θ*p. and were set as 1 × 10^−4^ and 5%, respectively.

### Microbiome profiling of external validation cohorts

A total of 1,716 publicly available gut metagenomic sequencing data were collected across five datasets^10,31–35^, including HMP (n=888), PRJEB15371 (n=55), PRJNA1086048 (n=601), PRJNA400072 (n=132), and PRJNA429990 (n=40). The quality control and host contamination removal were performed by KneadData. Species-level abundance profiles were generated using Phanta^56^, a tool designed for fast and accurate quantification of relative abundances of both prokaryotes and viruses. While our ORCA cohort was profiled using MetaPhlAn4, the validation cohorts—shared from an ongoing study by a co-author—were processed using Phanta and used solely to validate key findings rather than for direct comparison.

### Statistical analysis

Microbial diversity measures were calculated at the species level using the ‘vegan’ R package. The alpha diversity was measured by the Shannon index. The beta diversity for community composition was measured by the robust Aitchison distance in the PCoA. The differences in microbiome composition by disease status were tested with the PERMANOVA using the ‘adonis’ function in the ‘vegan’ R package. All PERMANOVA tests were performed with the 9,999 permutations based on the robust Aitchison distance.

In analyses testing our primary research questions, we used multivariate linear-mixed models in MaAsLin2^28^ (microbiome multivariable associations with linear models) to evaluate associations of microbial species and pathways with host factors while adjusting for the covariates, including age, gender, and smoking. Nominal p values across all associations were then adjusted for multiple comparisons using the Benjamini–Hochberg method with a target rate of 0.25 for *q* values^57^. Spearman correlations between metabolites and bacterial species were computed using the ‘corr.test’ function from the ‘psych’ R package. All statistical analyses were performed in R (version 4.4.0).

## Data availability

The metagenomic data from this study will be released upon acceptance.

## Code availability

The codes for statistical analyses and visualization are available in the GitHub repository (https://github.com/KelabatOSU/IBD_microbiome).

## Acknowledgements

This work was supported by the Resnek Family Center for PSC Research. We thank Dr. Chen Liao for useful discussions on the translocation analysis.

## Contributions

J.R.K. and Y.-Y.L. conceived and designed the project. S.K. and Z.Z. performed all the data analysis. S.K., J.R.K., and Y.-Y.L., interpreted the results and prepared the manuscript. X.Y., Y.Y., and Z.S. reviewed and edited the manuscript. M.D., K.W., and S.D.M. collected biological samples. All authors approved the manuscript.

