## Supplementary Information for "Oral–Gut Microbial Axis in Inflammatory Bowel Disease and Primary Sclerosing Cholangitis"

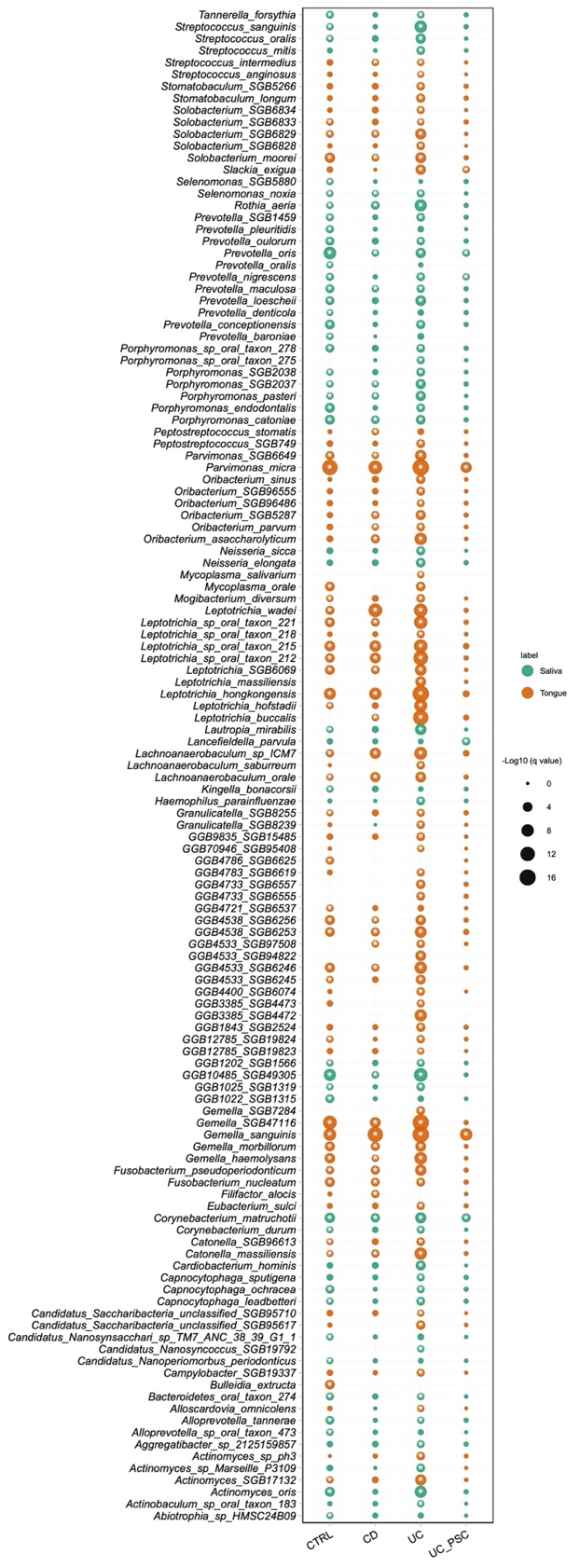

**Fig. S1. The species differences between saliva and tongue swabs in different phenotypic groups were identified using MaAsLin2.** Only statistically significant associations with q value ≤ 0.25 (Benjamini Hochberg adjusted P value) are labeled with an asterisk. The size of each dot represents the -log10 (q value). The color of each dot represents the valence of the association: green, enriched in saliva. Orange, enriched in tongue.

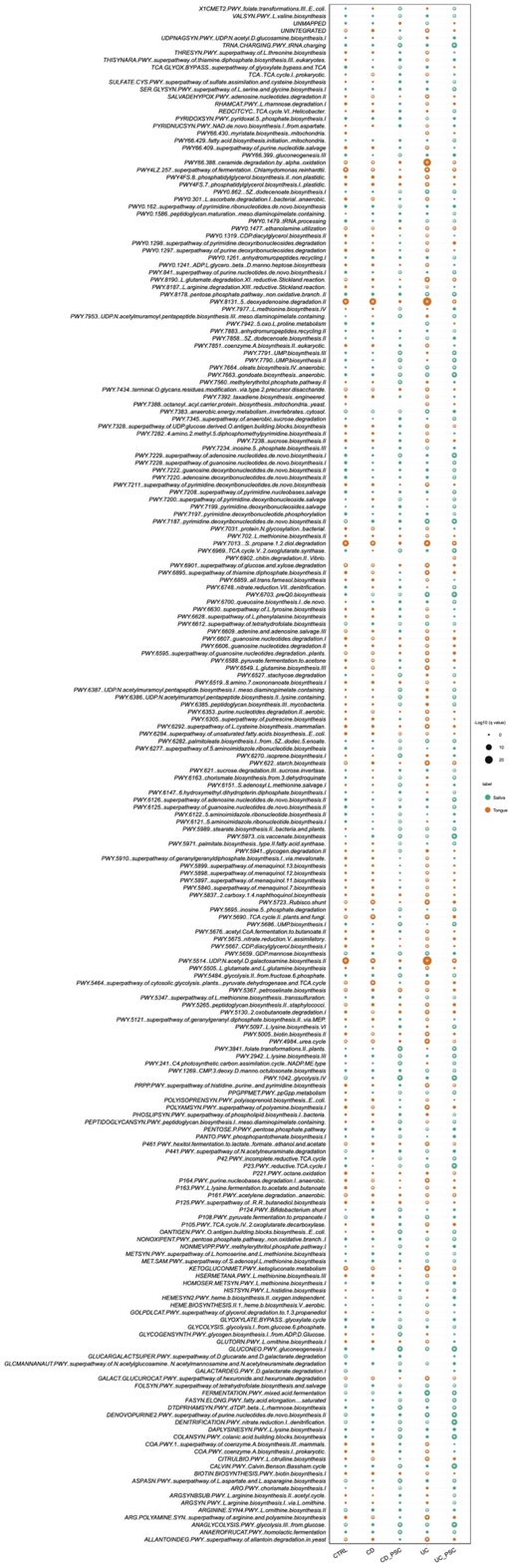

**Fig. S2. The functional capacity differences between saliva and tongue swabs in different phenotypic groups** **were** **identified using MaAsLin2.** Only statistically significant associations with q value ≤ 0.25 (Benjamini Hochberg adjusted P value) are labeled with an asterisk. The size of each dot represents the -log10 (q value). The color of each dot represents the valence of the association: green, enriched in saliva. Orange, enriched in tongue.

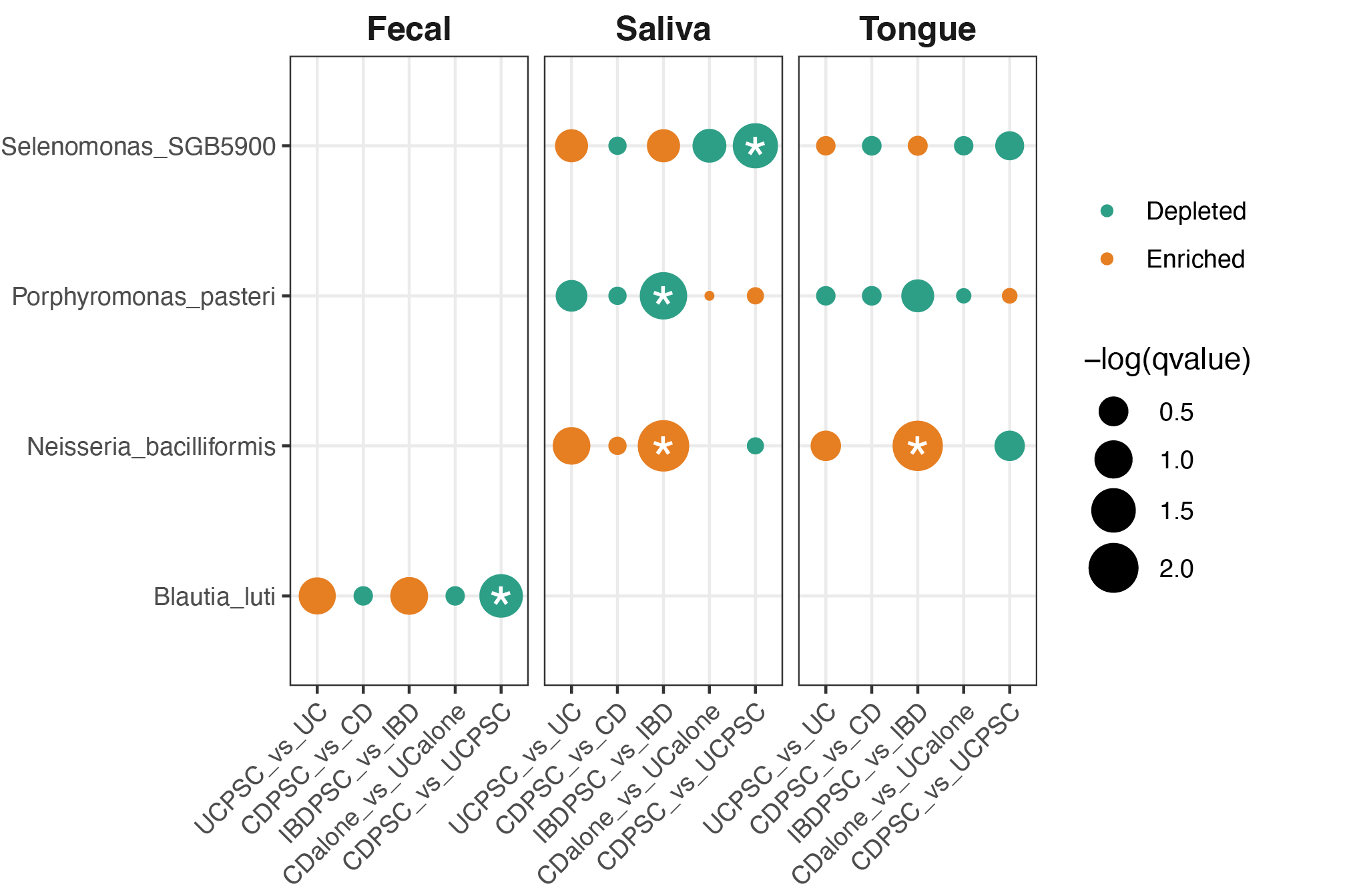

**Fig. S3. The species differences between disease groups (UC-PSC vs UC, CD-PSC vs CD, IBD-PSB vs IBD, CD alone vs UC alone, CD-PSC vs UC-PSC) across different body sites were identified using MaAsLin2.** Only statistically significant associations with q value ≤ 0.25 (Benjamini Hochberg adjusted P value) are labeled with an asterisk. The size of each dot represents the -log10 (q value). The color of each dot represents the valence of the association (green, depleted in cases; orange, enriched in cases). All comparisons are presented in a case-versus-control format (case_vs_control).

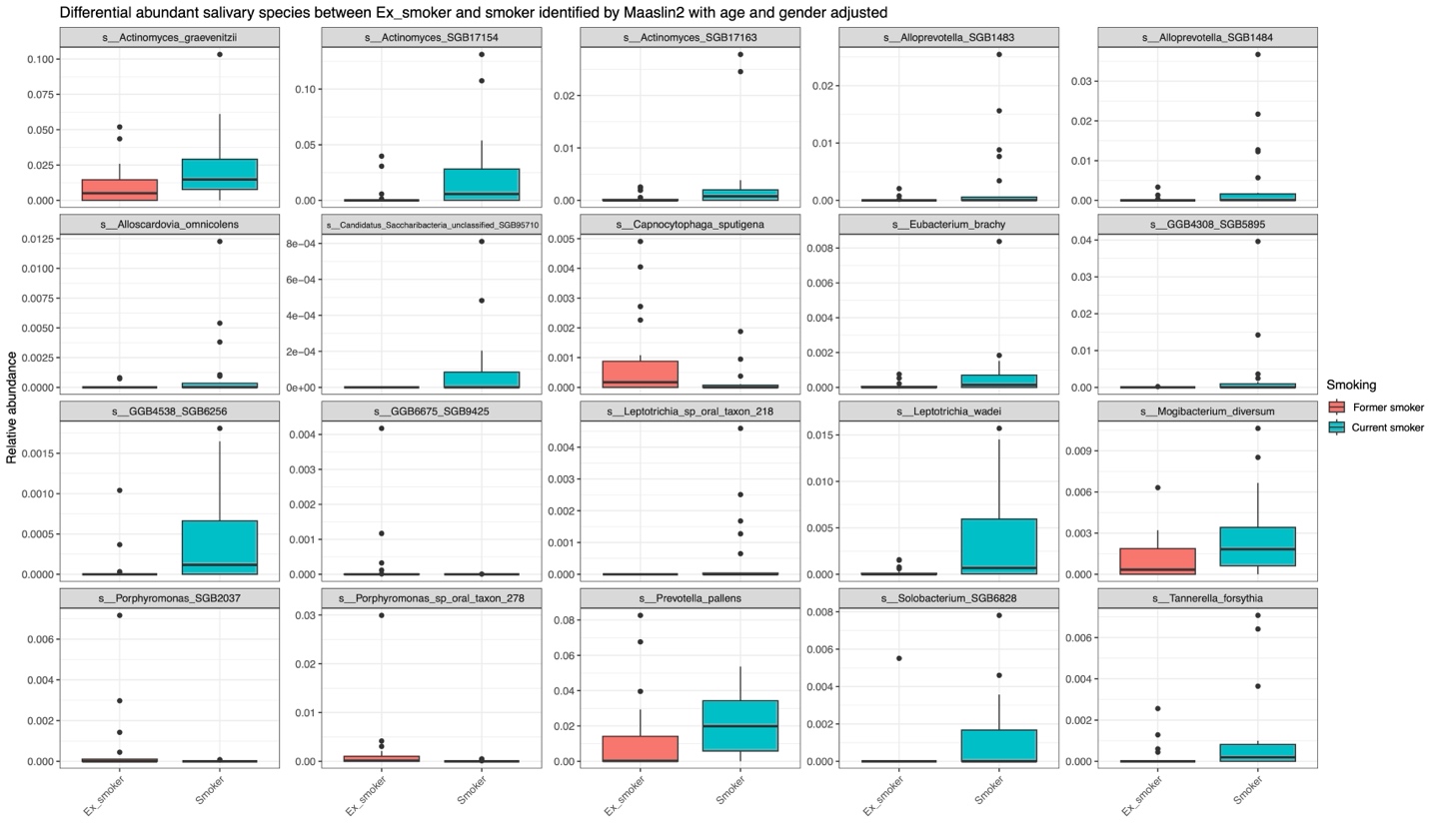

**Fig. S4. The salivary species differences between current smokers and former smokers in controls** were **identified using MaAsLin2.** Only statistically significant associations with q value ≤ 0.25 (Benjamini Hochberg adjusted P value) were plotted.

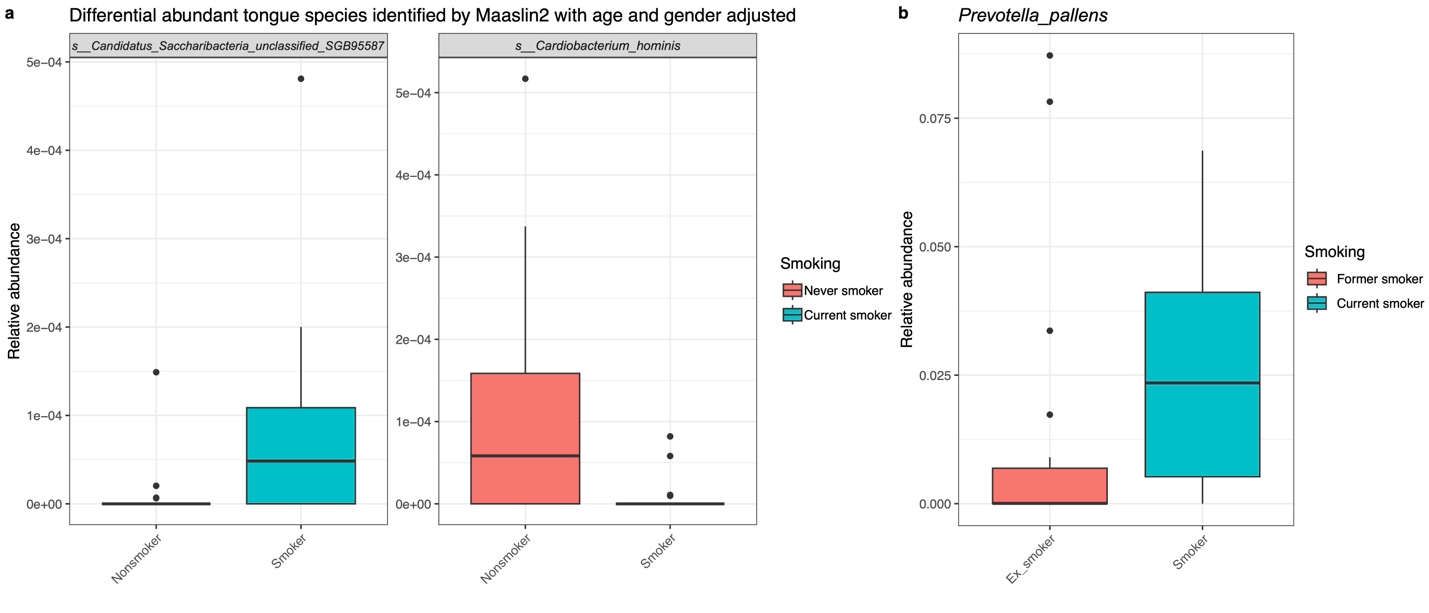

**Fig. S5. Smoking-related species features in tongue swab samples of healthy controls** **identified using MaAsLin2.** **a**, Comparison between current smokers and never smokers. **b**, Comparison between current smokers and former smokers. Only statistically significant associations with q value ≤ 0.25 (Benjamini Hochberg adjusted P value) were plotted.

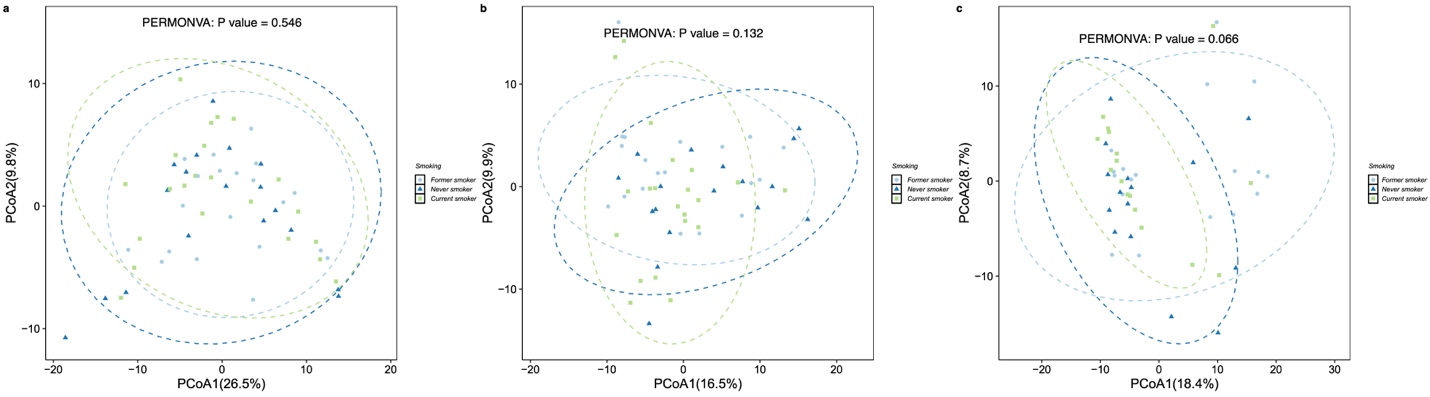

**Fig. S6. PCoA plot of microbiome samples from different smoking statuses at the functional capacity level based on the robust Aitchison distance. a**, Saliva. **b**, Tongue swap. **c**, Feces.

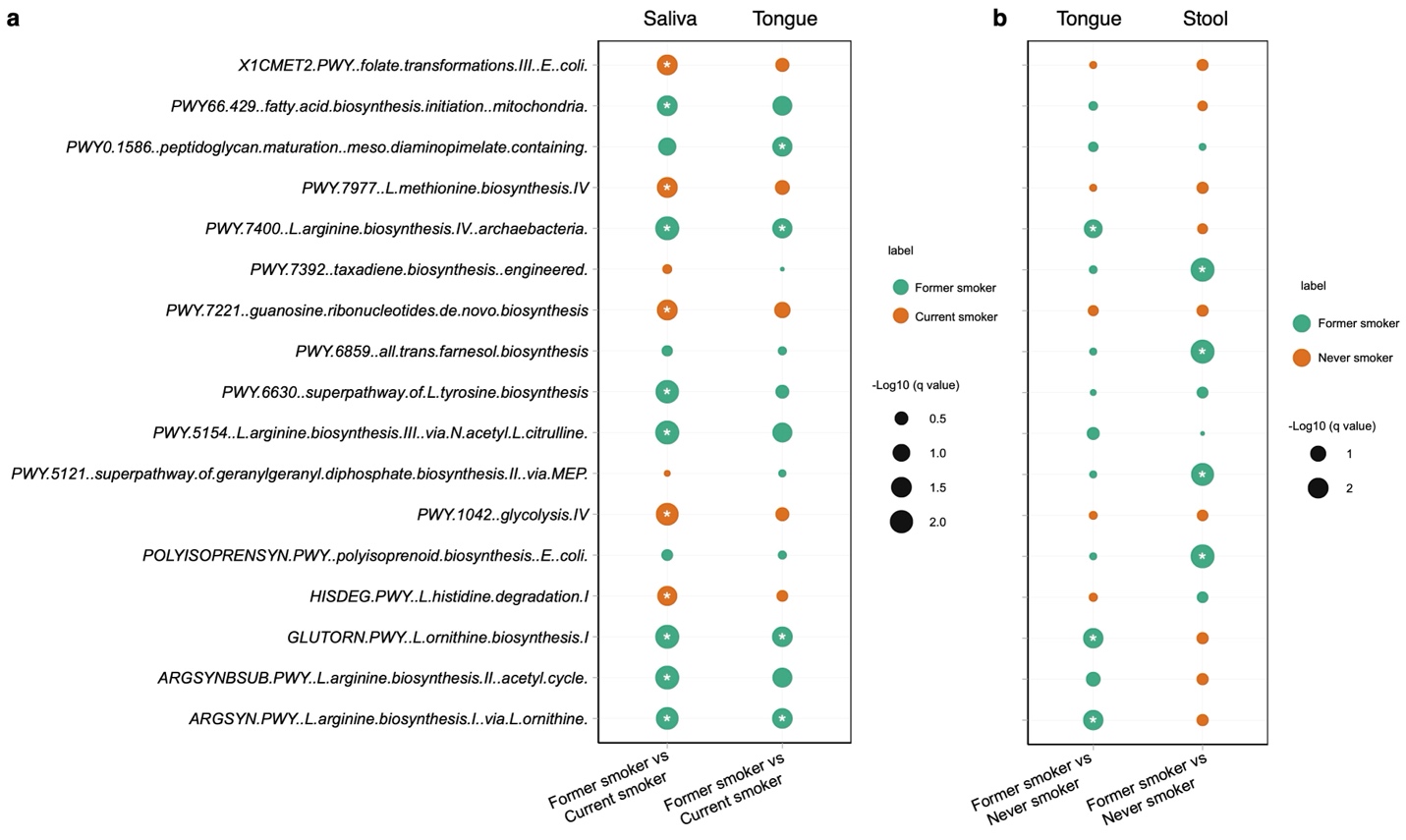

**Fig. S7** **Smoking-related functional pathways identified using MaAsLin2**. Significant smoking-related pathway features in saliva (**a**) and tongue swab (**b**) from each group are summarized. Only statistically significant associations with q value ≤ 0.25 (Benjamini Hochberg adjusted P value) are labeled with an asterisk. The size of each dot represents the log10 (q value). The color of each dot represents the valence of the association.

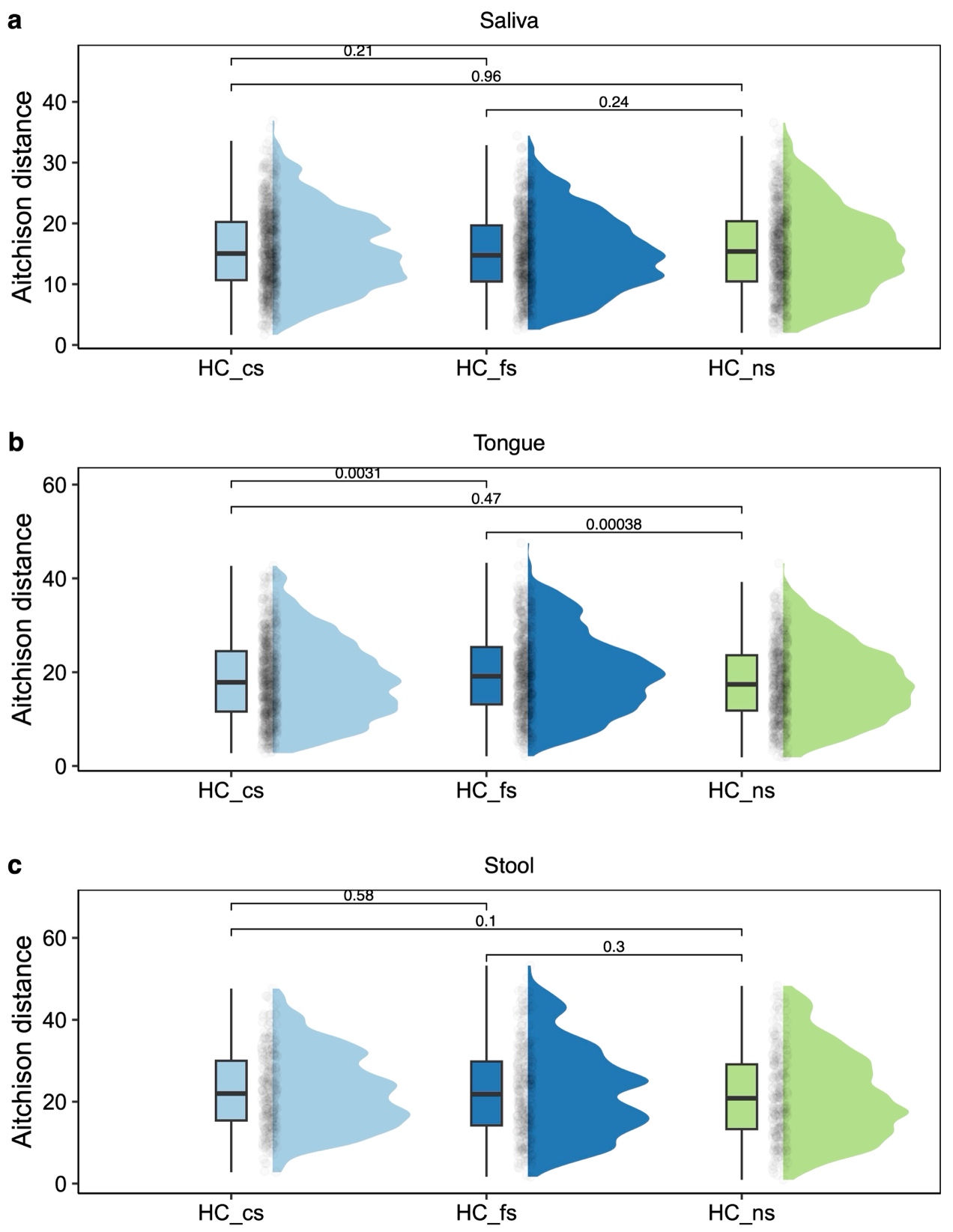

**Fig. S8 Associations between smoking history and oral microbiome in participants with CD.** Microbial differences between the smoking group and participants with CD quantified by robust Aitchison distance in saliva (**a**), tongue swab (**b**), and stool (**c**) samples. *P*-values were calculated by two-sided Mann–Whitney test. HC_ns_: Never smoker in Control group; HC_fs_: Former smoker in Control group; HC_cs_: Current smoker in Control group.
